# Differential oxidative costs of locomotory and genital damage in an orb-weaving spider

**DOI:** 10.1101/867325

**Authors:** Pierick Mouginot, Gabriele Uhl, Nia Toshkova, Michaël Beaulieu

**Affiliations:** Zoological Institute and Museum, University of Greifswald, Greifswald 17489, Germany; National Museum of Natural History, Sofia 1000, Bulgaria

**Keywords:** wound, tissue loss, genital damage, harmful male trait, oxidative status, *Larinia jeskovi*

## Abstract

In animals that regularly experience tissue loss, physiological responses may have evolved to overcome the related costs. Changes in oxidative status may reflect such self-maintenance mechanisms. Here, we investigated how markers of oxidative status varied in female orb-weaving spiders (*Larinia jeskovi*) by mimicking two distinct types of tissue loss they may naturally encounter: damage to their locomotory system and damage to their external genital structure, as inflicted by males to females during copulation (external female genital mutilation). Damage to the locomotory system resulted in a significant shift in the oxidative status reflecting investment into self-maintenance. In contrast, the loss of the genital structure did not result in quantitative changes of oxidative markers. The lack of response to genital mutilation suggests that genital mutilation is physiologically not costly for female spiders. The cost incurred to females rather arises from genital mutilation preventing the females from remating with another male.

## 1. Introduction

In nature, tissue loss typically occurs because of sub-lethal predation, agonistic behaviours between conspecifics and abiotic physical damage [1]. Tissue loss may be costly when it decreases the overall performance of injured animals by limiting their ability to exploit resources (*e.g.* reduced locomotor or feeding ability) and by affecting their homeostasis (*e.g.* because of fluid loss or infection). By activating physiological processes such as healing and immune responses, injured animals may minimize these direct costs. However, investing in these self-maintenance mechanisms may limit resources available for other fitness-related functions, such as reproduction [2]. This investment trade-off may be mediated by variation in oxidative status. Antioxidant defences can neutralize the action of oxidizing species on biomolecules, thereby limiting the generation of oxidative damage in tissues and increasing the survival probability of the organism [3]. For instance, *Bicyclus anynana* butterflies solve the trade-off between longevity and fecundity under challenging conditions by increasing antioxidant defences, thereby prolonging lifespan but reducing fecundity [4]. Alternatively, to facilitate a healing and immune response following tissue loss, injured organisms may also locally reduce their antioxidant response, as oxidizing species may themselves enhance cell communication during tissue repair and eliminate pathogens [5, 6]. Moreover, in order to minimize oxidative damage on their own tissues, injured animals may simultaneously reduce their physical activity thereby reducing their overall production of oxidizing molecules [4]. Hence, the optimal maintenance response of injured animals depends on the regulation of the balance between their production of oxidizing molecules and their antioxidant response [7].

Tissue loss does not only occur because of predation or agonistic behaviours between conspecifics but also during copulation. Indeed, in a broad range of species, males can harm females while transferring sperm and seminal fluids by inflicting physical damage inside or outside females’ genitalia [8]. Because sperm and seminal fluids represent resources that females may use, they can affect the physiology of inseminated females [8, 9] and may therefore affect the physiological response of females following damage during copulation. In several spider species, males mutilate the outer structures (scapus) of the female genitalia [10, 11], which makes it possible to disentangle the actual costs due to genital damage from other copulation effects. The physiological response of females following genital damage has previously been investigated with treatments mimicking genital damage by ablating a locomotory tissue [12]. However, locomotory and genital damage may not elicit a similar response. Understanding this response would clarify the mechanisms underlying sexual conflict in species where female genital mutilation occurs.

Here, we investigated the physiological response to locomotory and genital damage by measuring different markers of oxidative status in female spiders experiencing tissue loss. To mimic locomotory and genital damage, we applied experimental ablation of one leg and of the scapus. If males and females coevolved to reduce the physiological costs of genital mutilation [12], we expected genital mutilation to trigger a weaker physiological response than locomotory tissue loss.

## 2. Material and Methods

### Study animals

We collected sub-adult (one moulting stage from adulthood) females of the orb-weaving spider *Larinia jeskovi*, Marusik 1986 (Araneidae) [13] in August 2015 and 2016 in the Biebrza National Park, Poland (53°21’01.36’’N, 22°34’37.45’’E). In the laboratory, we housed females individually in 250 mL plastic cups at room temperature and under a natural light cycle. Females were fed with one fly (*Lucilia sericata*) every three days and watered daily. Sub-adult females (one moulting stage from adulthood) were checked daily for moulting events. After their final moult, adult females were used for the experimental setup.

### Experimental setup (figure 1)

In order to assess the effect of tissue loss on females’ oxidative status, we amputated part of one of their forelegs (randomly right or left leg) or manually ablated their scapus [10]. To achieve this, females were first immobilized under a mesh and then wounded under a stereomicroscope. A control group was left intact but was similarly handled. In 2015, 30 females were randomly assigned to three treatment groups: control, tibia-amputated, scapus-amputated. In 2016, we repeated the procedure with an additional wounding treatment in which we removed 200 μm of the tarsus. The removal of only the tip of the leg was chosen to assess the effect of a wound comparable to the scapus mutilation in terms of the amount of tissue lost. Seventy-two females were randomly assigned to the four experimental amputation treatments; control, mid-tibia, leg-tip and scapus. Eight hours after treatment, females were cryofixed with liquid nitrogen and stored at −80 °C for later analysis. Age was calculated as the number of days between the final moult and the treatment.

**Figure 1:**
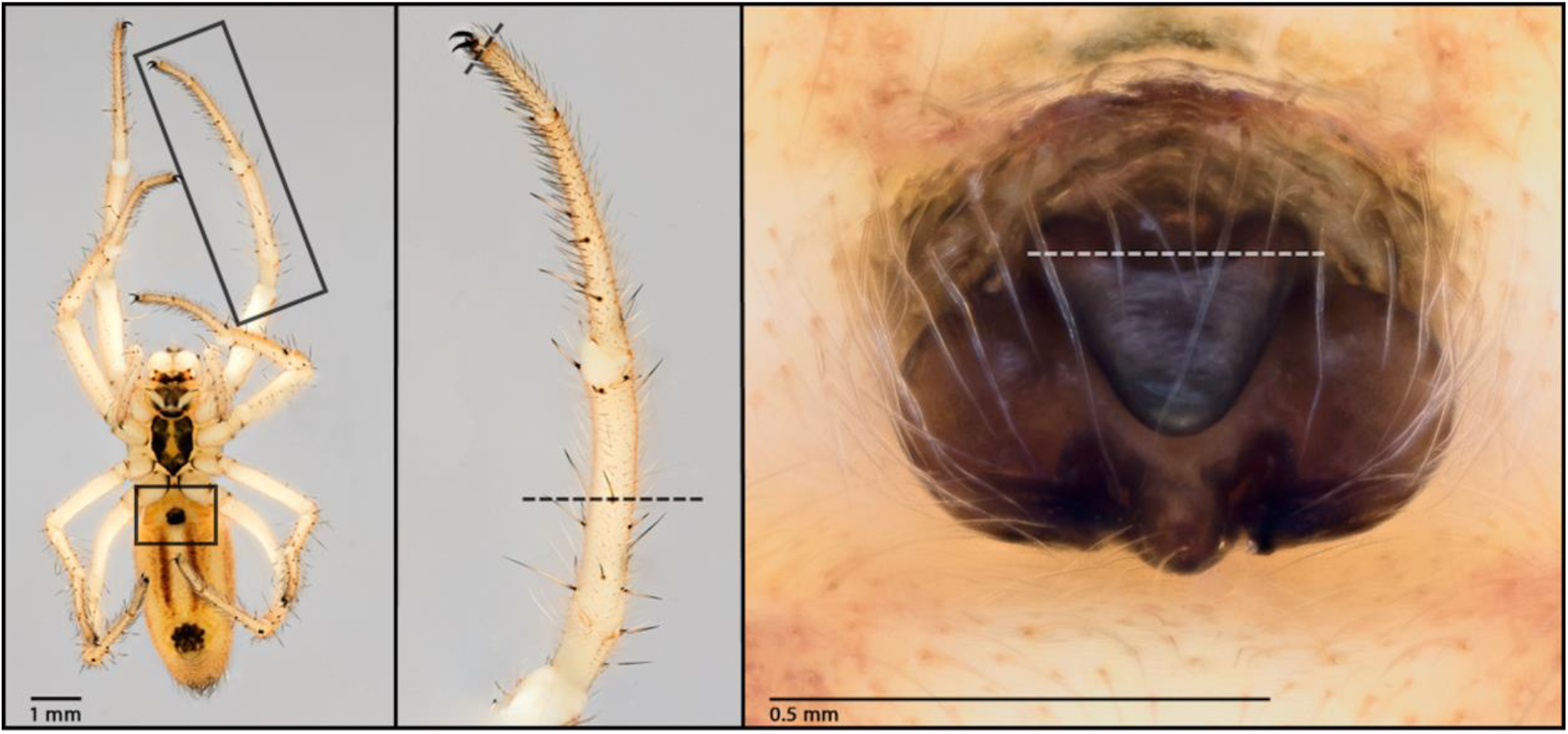
*Larinia jeskovi* female in ventral view (A). The two black rectangles indicate the leg and the genital areas. Close up picture of the leg with the experimental amputation treatments at the mid-tibia and leg-tip represented by the dashed lines (B). Close up picture of the external female genitalia with the experimental amputation treatment applied at the basis of the scapus represented by the dashed line (C).

### Oxidative stress markers

Before measurements, all appendages (legs, pedipalps) were removed from the frozen specimens on dry ice, and the body of each female was weighed to the nearest 0.01 mg (Sartorius LE225D; Sartorius AG, Göttingen, Germany) before being mixed with PBS buffer.

In 2015, we measured two markers of oxidative status. We used the OXY-absorbent test (Diacron International, Grosseto, Italy) to measure spiders’ total antioxidant capacity (expressed in millimole of HOCL neutralized) and the d-ROM test (Diacron International, Grosseto, Italy) to measure concentrations in hydroperoxides, a marker of oxidative damage deriving from the oxidation of fatty acids, proteins and nucleic acids and promoting cell death (expressed in milligrams per decilitre of H_2_O_2_ equivalent). For both tests, we followed the procedure described in [14]. Three individuals, for which d-ROM values were below the detection threshold, were excluded from the analyses. This resulted in a sample size of 9 control females, 10 tibia-amputated females, and 8 scapus-amputated females.

In 2016, we measured glutathione (GSH) levels (expressed in μmol GSH per mg protein) as marker of endogenous antioxidant defences, and malondialdehyde (MDA) levels to assess oxidative damage on lipids (expressed in mmol per mg protein). The thorax and the abdomen were homogenized together with Triton buffer (7.5 μl for each 1 mg sample) through high-speed shaking (three times for 1 min; 24 shakes/s). The resulting homogenate was centrifuged (16249 g, 30 min, 4°C) before transferring the resulting supernatant to a new tube and centrifuging it again (16249 g, 15 min, 4°C). The second supernatant was then used to analyse total protein and MDA concentrations. MDA concentrations were determined using the commercial kit MDA Microplate Assay Kit (Cat. no. CAK1011; Cohesion Bioscience; 532 and 600 nm). GSH levels were assessed with spectrophotometric method, which involves oxidation of GSH by the sulfhydryl reagent 5,5′-dithio-bis (2-nitrobenzoic acid) (DTNB, also known as Ellman’s reagent) to form the yellow derivative 5′-thio-2-nitrobenzoic acid (TNB), measurable at 412 nm. Total protein concentration of all samples was determined using the Bradford protein assay at 595 nm. Two females died before the end of the treatment. Out of the 70 females left, we were able to measure MDA levels in 62 individuals (others showing levels lower than the minimal detection threshold). Consequently, 70 females were used for glutathione measurements (18 control females, 18 leg-tip-amputated females, 17 tibia-amputated females, and 17 scapus-amputated females) and 62 females were used for MDA measurements (17 control females, 15 mid-tibia-amputated females, 16 leg-tip-amputated females, and 14 scapus-amputated females).

### Statistical analyses

To test the effects of treatment on antioxidant capacity, hydroperoxide and MDA levels, we built linear models with antioxidant capacity, hydroperoxide or MDA levels as dependent variables, and treatment, age, body mass and protein concentration (to correct for concentration differences between samples) as independent variables. To test the effect of treatment on glutathione levels, we built a linear model with glutathione corrected for body mass as dependent variable (to reach normality) and treatment and age as independent variables. We checked linearity assumptions graphically as well as with a Bartlett test for homoscedasticity and Shapiro test for normality of the residuals. The model for antioxidant capacity required the exclusion of one outlier (from control treatment) to match linearity assumptions. There was no significant correlation among variables (table S1, Suppl. Mat.) leading to no collinearity among explanatory variables. In all models, quantitative variables were centred and standardized [15].

For each model, we considered all plausible candidate models and ranked them according to their AICc value [16, 17]. To evaluate the contribution of each predictor to the model prediction, we calculated its sum of Akaike weights and used “full model averaging” to calculate parameter estimates β [17]. Since the sum of weights may provide a poor evaluation of the predictors’ importance [18], we calculated the 85% confidence interval for each parameter estimate [19]. Parameter estimates whose confidence interval did not include zero were considered as having a significant effect. The evaluation of predictors’ contribution results in parameter estimates for which the first level of a factor is set as a reference. Thus, the results of the amputation treatments are presented as mid-tibia, leg-tip and scapus amputation treatments compared to the control treatment as the reference. All analyses were performed in R software [20]. The MuMIn v1.40.4 [21] and Plotrix 3.7-6 [22] packages were used for procedures of model selection and model averaging and for calculating the standard errors of the means in figure 2, respectively.

**Figure 2:**
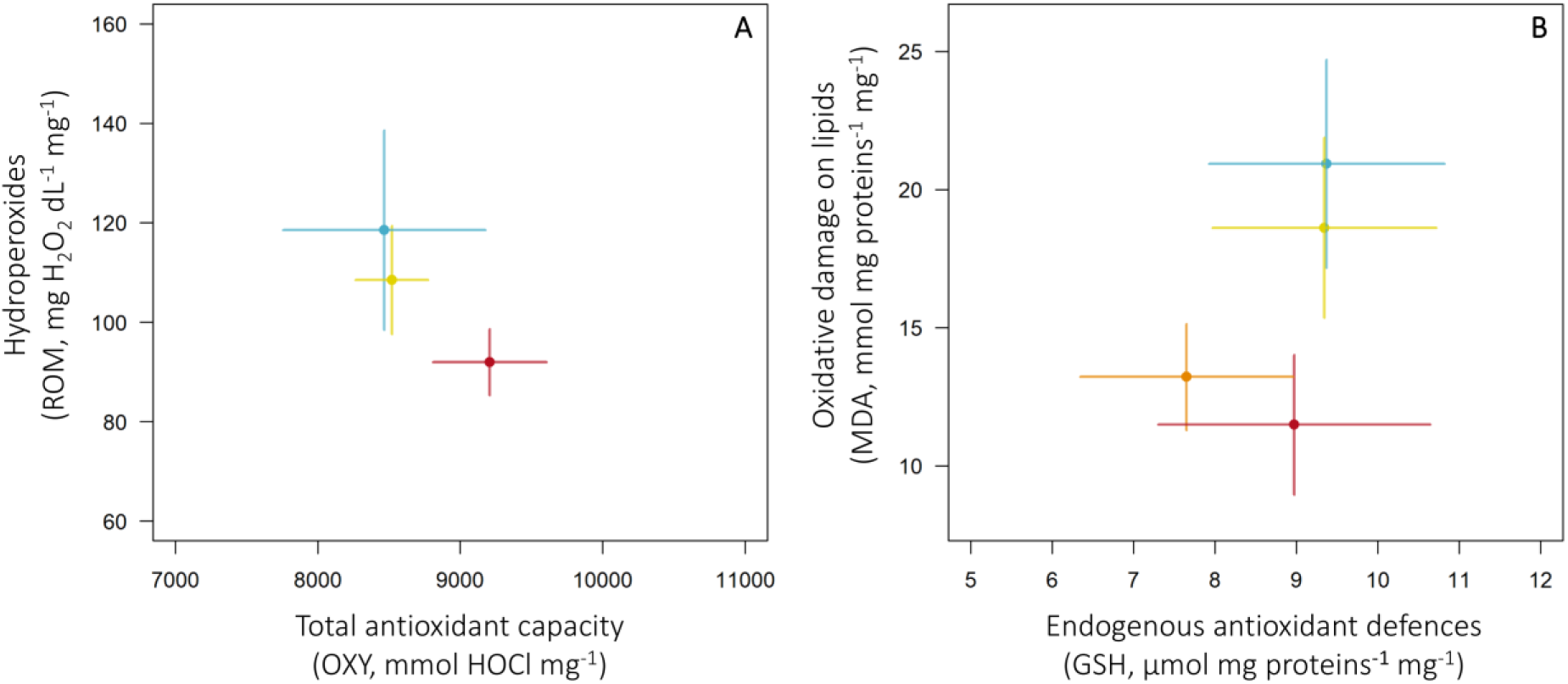
Mean values (± SE) of A: total antioxidant capacity and oxidative damage (hydroperoxide concentration), B: endogeneous antioxidant defence (glutathione concentration) and oxidative damage on lipids (MDA concentration) for each experimental amputation treatment: control (blue), mid-tibia (red), legtip (orange) and scapus (yellow) in the spider *Larinia jeskovi*. Markers of antioxidant defences are represented on the x-axes and markers of oxidative damage on the y-axes. Levels of antioxidant defences and oxidative damage markers are corrected for body mass.

## 3. Results

Spiders did not differ in body mass and age between treatments (Suppl. Mat., figure S1). In 2015, females amputated at the mid-tibia showed higher antioxidant capacity and lower hydroperoxide levels than control females (table 1). Similarly, in 2016, both mid-tibia and leg-tip amputations were associated with lower MDA levels compared to control females. However, glutathione levels were similar in all treatment groups (table 1). Scapus-amputated females did not show any alteration of their oxidative status relative to control females both in 2015 and 2016 (figure 2, table 1). As a result, leg-amputated spiders segregated from scape-amputated and control spiders in oxidative space (figure 2).

**Table 1:**
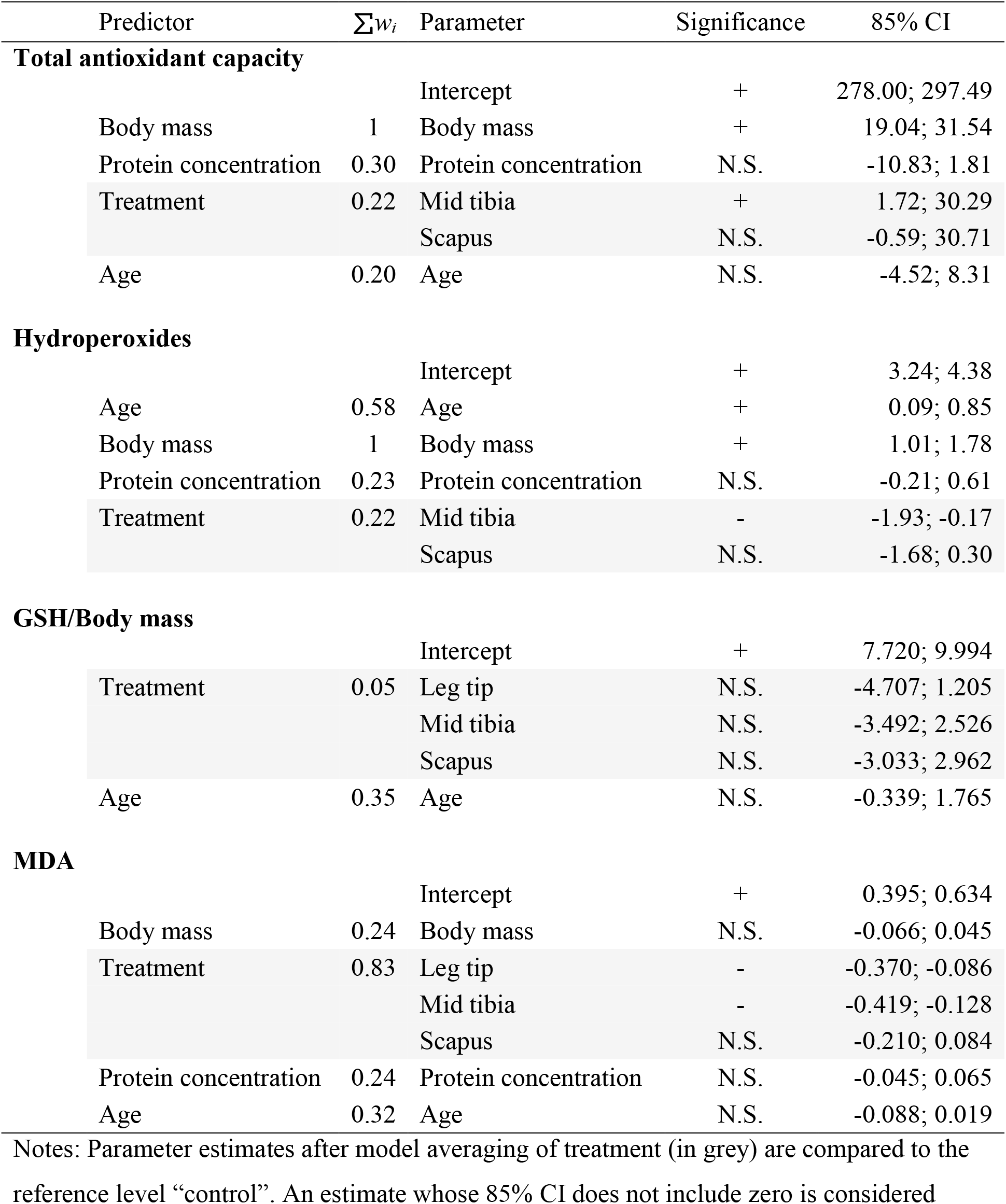
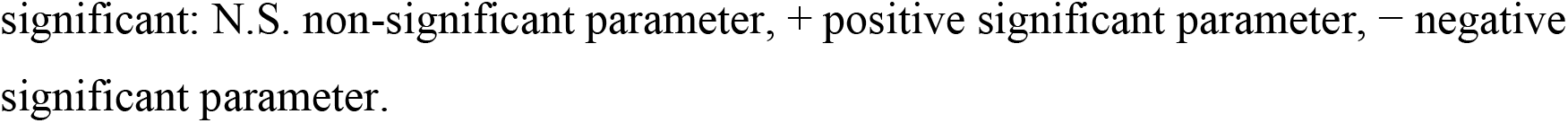
Predictor’s sum of weights, parameter significance and 85% confidence interval (CI) after full model averaging on the set of candidate models, assessing the effect of body mass, experimental amputation treatment, age and protein concentration on levels of antioxidant defence (estimated by the total antioxidant capacity test and GSH levels) and oxidative damage (estimated by hydroperoxide levels and MDA levels) in the spider *Larinia jeskovi*.

| Predictor | $\Sigma w_i$ | Parameter | Significance | 85% CI |
| --- | --- | --- | --- | --- |
| **Total antioxidant capacity** |  |  |  |  |
|  |  | Intercept | + | 278.00; 297.49 |
| Body mass | 1 | Body mass | + | 19.04; 31.54 |
| Protein concentration | 0.30 | Protein concentration | N.S. | -10.83; 1.81 |
| Treatment | 0.22 | Mid tibia | + | 1.72; 30.29 |
|  |  | Scapus | N.S. | -0.59; 30.71 |
| Age | 0.20 | Age | N.S. | -4.52; 8.31 |
| **Hydroperoxides** |  |  |  |  |
|  |  | Intercept | + | 3.24; 4.38 |
| Age | 0.58 | Age | + | 0.09; 0.85 |
| Body mass | 1 | Body mass | + | 1.01; 1.78 |
| Protein concentration | 0.23 | Protein concentration | N.S. | -0.21; 0.61 |
| Treatment | 0.22 | Mid tibia | - | -1.93; -0.17 |
|  |  | Scapus | N.S. | -1.68; 0.30 |
| **GSH/Body mass** |  |  |  |  |
|  |  | Intercept | + | 7.720; 9.994 |
| Treatment | 0.05 | Leg tip | N.S. | -4.707; 1.205 |
|  |  | Mid tibia | N.S. | -3.492; 2.526 |
|  |  | Scapus | N.S. | -3.033; 2.962 |
| Age | 0.35 | Age | N.S. | -0.339; 1.765 |
| **MDA** |  |  |  |  |
|  |  | Intercept | + | 0.395; 0.634 |
| Body mass | 0.24 | Body mass | N.S. | -0.066; 0.045 |
| Treatment | 0.83 | Leg tip | - | -0.370; -0.086 |
|  |  | Mid tibia | - | -0.419; -0.128 |
|  |  | Scapus | N.S. | -0.210; 0.084 |
| Protein concentration | 0.24 | Protein concentration | N.S. | -0.045; 0.065 |
| Age | 0.32 | Age | N.S. | -0.088; 0.019 |

Notes: Parameter estimates after model averaging of treatment (in grey) are compared to the reference level "control". An estimate whose 85% CI does not include zero is considered
significant: N.S. non-significant parameter, + positive significant parameter, – negative
significant parameter.

## 4. Discussion

Leg amputation led to a shift in the oxidative status of female spiders irrespective of amputation extent. In contrast, scapus amputation did not affect their oxidative status. These results were consistent across different markers of oxidative damage measured in different individuals in two distinct experiments. Hence, our study suggests that a physical harm inflicted to the locomotory system of female spiders affects their oxidative balance, whereas a damage to their external genitalia does not.

In agreement with our predictions, tissue loss due to leg amputation induced a shift in females’ oxidative balance, as amputated females showed higher antioxidant capacity than intact females. However, glutathione levels were similar in all treatment groups. This pattern suggests that amputated females invested in self-maintenance mechanisms by upregulating their production of some endogenous antioxidant defences, which did not involve glutathione or involved glutathione in combination with other antioxidant compounds. This upregulation was not associated with stable oxidative damage in amputated females but with lower oxidative damage. Low oxidative damage in amputated females is likely related to their probable lower locomotory activity following leg amputation leading to lower production of oxidizing molecules [4]. Moreover, low physical activity in amputated females may help them to save resources that can be allocated to maintenance mechanisms, such as antioxidant defences. Finally, the fact that both markers of oxidative damage decreased in amputated individuals can be explained by the biochemical proximity between both markers, as MDA results from the decomposition of hydroperoxides [23].

In contrast to leg amputation, the removal of external genital structure did not affect the females’ oxidative status, which was comparable to that of intact females. It seems possible that the extent of the harm was not sufficient to induce a detectable physiological response. However, the amputation of the leg tip, an injury comparable to scapus mutilation in terms of tissue loss, affected the females’ oxidative balance. This difference may be due to the negative effect of locomotory tissue loss on the overall physical activity of spiders, which is unlikely to occur following genital mutilation [4]. Since locomotory and genital damage trigger different physiological responses, we argue that ablating a locomotory tissue to mimic the effects of genital damage is not relevant [12], and only genitally-mutilated females should be used to examine the effects of such mutilation.

External female genital mutilation by males is a common feature of the mating system of several spider species that allows males to secure paternity [10, 11]. As a harmful male adaptation, the mutilation of female genitalia may be subject to a sexual conflict [8]. However, the costs and benefits for females are unclear. Our findings suggest that female spiders do not experience oxidative changes due to genital mutilation, suggesting no costs in terms of self-maintenance. The absence of physiological costs might therefore be the result of selection on males and females to reduce the costs associated with a harm that would ultimately result in lower reproductive performance in both females and males [9, 12, 24]. Our results therefore corroborate the oxidative shielding hypothesis, postulating that oxidative damage in reproductive females is minimized to avoid deleterious effects on reproductive performance [25]. Overall, our study suggests that males benefit from mutilating female genitalia without impairing female’s fecundity, which sheds new light on the evolution of external female genital mutilation in animals [26].

## Ethical statement

All spiders were handled with care and cold-euthanized.

## Data accessibility

Data are available from the Dryad Digital Repository: [27]

## Author contributions

P.M., G.U. and M.B. designed the study and contributed through discussions. P.M. collected animals and carried out experimental treatments. P.M. and N.T. carried out physiological measurements. P.M. and M.B. carried out statistical analyses. All authors contributed through writing.

## Competing interests

No competing interests declared.

## Funding

P.M. was funded by the German Science Foundation (Uh87/7-1 to GU). N.T. was supported by an exchange grant of the research training group RESPONSE funded by the German Research Foundation (DFG GRK2010). We thank Klaus Fischer for supporting the study.

## Acknowledgements

We thank P.O.M. Steinhoff and R. Fragueira for help during experiments and physiological measurements. We are grateful to P. Michalik for providing pictures (Visionary Digital BK Plus Lab System) and Janusz Kupryjanowicz for logistic support.

